# Dynamics of a mutual inhibition between pyramidal neurons compared to human perceptual competition

**DOI:** 10.1101/2020.05.26.113324

**Authors:** N. Kogo, F. B. Kern, T. Nowotny, R. van Ee, R. van Wezel, T. Aihara

## Abstract

Neural competition plays an essential role in active selection processes of noisy and ambiguous input signals and it is assumed to underlie emergent properties of brain functioning such as perceptual organization and decision making. Despite ample theoretical research on neural competition, experimental tools to allow neurophysiological investigation of competing neurons have not been available. We developed a “hybrid” system where real-life neurons and a computer-simulated neural circuit interacted. It enabled us to construct a mutual inhibition circuit between two real life pyramidal neurons. We then asked what dynamics this minimal unit of neural competition exhibits and compared them to the known behavioral-level dynamics of neural competition. We found that the pair of neurons shows bi-stability when activated simultaneously by current injections. The addition of modelled noise and changes in the activation strength showed that the dynamics of the circuit are strikingly similar to the known properties of bi-stable visual perception.

## Introduction

Visual perception is an emergent property resulting from an active organization of input signals by the brain. This organization has to be accomplished while being subjected to the underrepresented, noisy and ambiguous signals received by the eyes. In other words, the brain is constantly challenged to make coherent selections among often conflicting local signals. Underlying the selection processes are neural competition mechanisms between neurons representing the conflicting signals. A well-known perceptual phenomenon representing signal competition and selection processes is “bi-stable perception” that occurs when visual signals support two likely perceptual interpretations. Signals that support one of the percepts are selected coherently at any given time and one percept becomes dominant. The input signals are eventually re-organized to establish the alternative percept, leading to reversals between the two percepts every few seconds (Leopold & Logothetis, 1999). This repetitive perceptual re-organization in bi-stable perception provides information about how visual signals are processed, organized, and eventually lead to conscious perception. The abundant literature on bi-stable perception is an important resource of information to investigate the neural mechanisms of signal competition and selection processes. In the computational neuroscience literature, neural competition is often modelled by “mutual inhibition” between differently tuned neurons. A possible neural circuit diagram of mutual inhibition is shown in Fig. 1a in which each pyramidal neuron (PN1 or PN2) activates a partner inhibitory neuron (IN1 and IN2, respectively) which, in turn, projects an inhibitory synapse to the competing pyramidal neuron, forming disynaptic inhibitory connections in both directions.

**Fig. 1.**
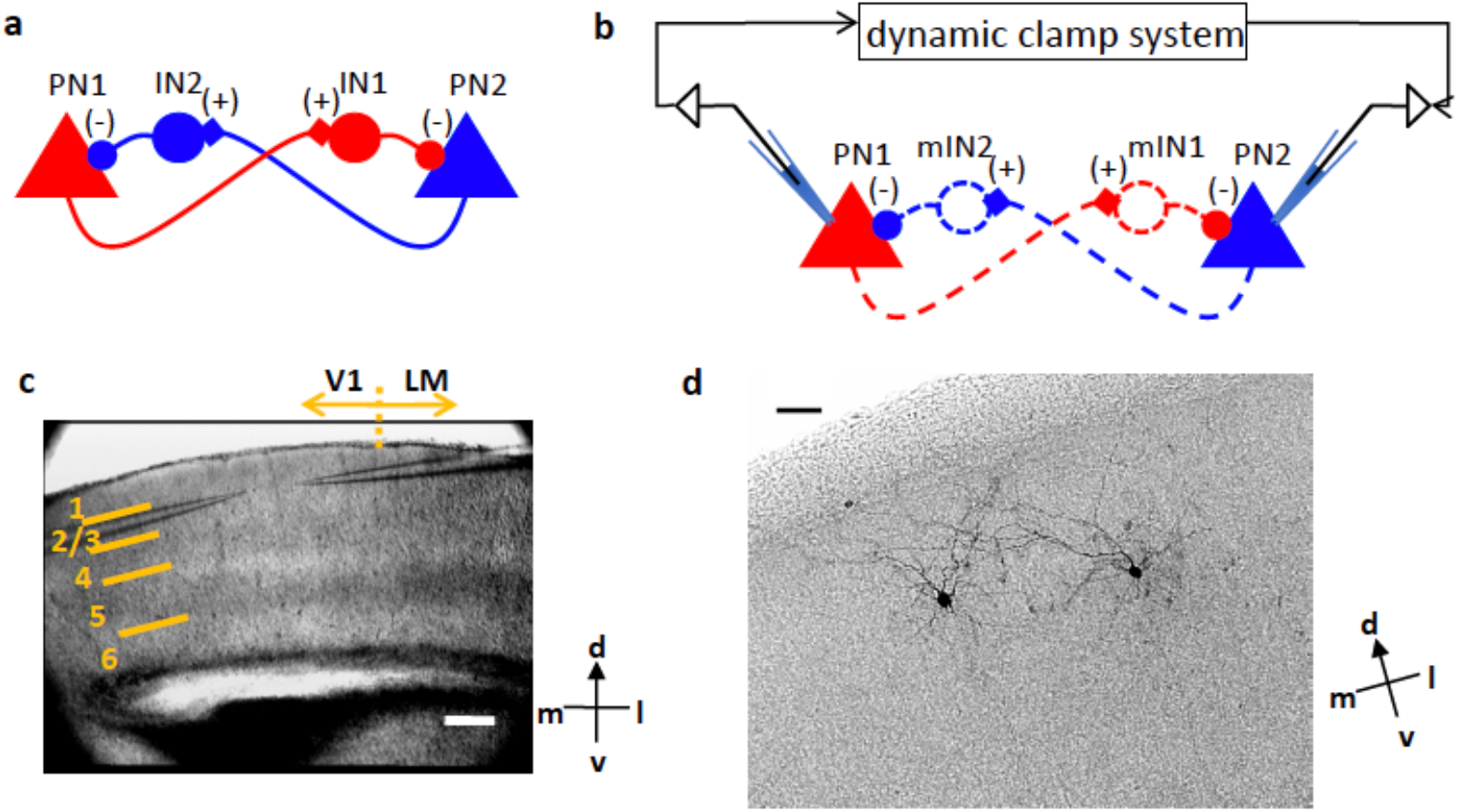
Mutual inhibition circuit and experimental design. **a**: Neural circuit diagram for a mutual inhibition. Triangles: pyramidal neurons (PNs). Disks: inhibitory neurons (INs). **b**: The disynaptic mutual inhibition circuit was established between two real-life pyramidal neurons by implementing model inhibitory neurons and synapses (dashed lines) in the StdpC dynamic clamp system. **c**: An image of the brain slice (right hemisphere) from a DIC-IR microscope during recording with two patch recording pipettes placed in layer 2/3 of V1. 1 to 6: six layers. LM: lateromedial area. d: dorsal, v: ventral, l: lateral, m: medial. Scale bar: 200 μm. **d**: An example of biocytin filled pair of pyramidal neurons. Scale bar: 50 μm.

It has been suggested that the conflicting signals for local features such as orientation (Bonds, 1989; Sillito, 1975), motion direction (Mikami et al., 1986; Snowden et al., 1991), and edge assignment (Kogo & van Ee, 2015; Zhou et al., 2000) compete with each other through such mutual inhibition circuits. This mutual inhibition circuit has been implemented in computer models to explain bi-stable perception (Laing & Chow, 2002; Lankheet, 2006; Matsuoka, 1984; Mueller, 1990; Noest et al., 2007; Shpiro et al., 2009; Wilson et al., 2000; Wilson, 1999), object recognition (Masquelier et al., 2009), decision making (Heuer, 1987; Machens et al., 2005; Usher & McClelland, 2001), and place cell field generation (Mark et al., 2017). It has also been suggested that these circuits underlie mechanisms such as larger scale neural interactions and feedback systems (Beck & Kastner, 2005; Lee et al., 1999; Wang et al., 2013) that establish a globally coherent percept. Moreover, disynaptic inhibitory connections between pyramidal neurons, the necessary constituent of mutual inhibition, are found in various layers and areas of neocortex (Berger et al., 2009; Kapfer et al., 2007; Ren et al., 2007; Silberberg & Markram, 2007) and hippocampus (Miles, 1990). It is hence possible that mutual inhibition serves as a canonical element of signal processing circuits in the brain.

Despite the numerous theoretical models implementing mutual inhibition circuits, experimental tools are missing that allow thorough neurophysiological analysis of competing cortical neurons at the system-wide level, due to the limitations of current technology. However, with the approach introduced in this paper, it is possible to construct a minimal unit of neural competition in real-life. By investigating the neural dynamics of the minimal unit, considering it as a building block of the whole system, and comparing its dynamics to the ones of the whole system, it may be possible to deduce how neural elements are integrated into a whole system such that known behavioral properties emerge.

We established an experimental model where a model mutual inhibition circuit is implemented between a pair of two real-life pyramidal neurons in brain slice preparations of mouse primary visual cortex (Fig. 1). The two neurons are patch clamped and connected with each other via a computer model that allows them to interact in real time. This hybrid system has the advantage of keeping all physiological properties of the real pyramidal neurons intact, while providing full control over the computer simulated connections between them. Using this hybrid system, we succeeded to evoke bi-stable activity in the pyramidal neurons. We investigated the dynamics of the bi-stable activity and compared them with the known dynamics of bi-stable visual perception, namely the effects of noise and the effect of changing stimulus input intensity.

## Results

Double patch clamp recordings were performed from visually identified pyramidal neurons in layer 2/3 of mouse primary visual cortex (Fig. 1c). In total, 93 pairs of pyramidal neurons from 32 mice were recorded. By using biocytin-filled patch pipettes, some pyramidal neuron pairs were labeled and visualized after the experiments (N=9). In all cases, the stereotypical morphology of pyramidal neurons (with a short apical dendrite and thin multiple oblique dendrites) was identified, located in layer 2/3 of V1 (Fig. 1d).

### Disynaptic mutual inhibitory connections

Mutual inhibitory connections between each pair of pyramidal neurons were constructed by a dynamic clamp system (spike timing dependent plasticity clamp, StdpC (Kemenes et al., 2011; Nowotny et al., 2006). The inhibitory neurons were modeled by implementing Hodgkin-Huxley type conductance-based ion channel models with parameters derived from literature (see Materials and Methods). The connections between the (real) pyramidal neurons and the (model) inhibitory neurons were established with modelled excitatory and inhibitory synapses (see Materials and Methods for details). In Fig. 2a, an example of the recording of two pyramidal neurons is shown. An action potential of pyramidal neuron 1 (PN1) was evoked by the injection of a short depolarization current pulse (red triangle). This action potential triggered the computation within the dynamic clamp system, activating a model inhibitory neuron, and, in turn, triggering an inhibitory synaptic conductance and current, which was injected into pyramidal neuron 2 (PN2) (see Fig. 7 for more details). This evokes an inhibitory postsynaptic potential (IPSP) in PN2 (blue asterisk in Fig. 2a). When an action potential was evoked in PN2 (blue triangle), a modelled IPSP was evoked in PN1 (red asterisk), illustrating that a mutual inhibition circuit between the pair of pyramidal neurons was established successfully.

**Fig. 2.**
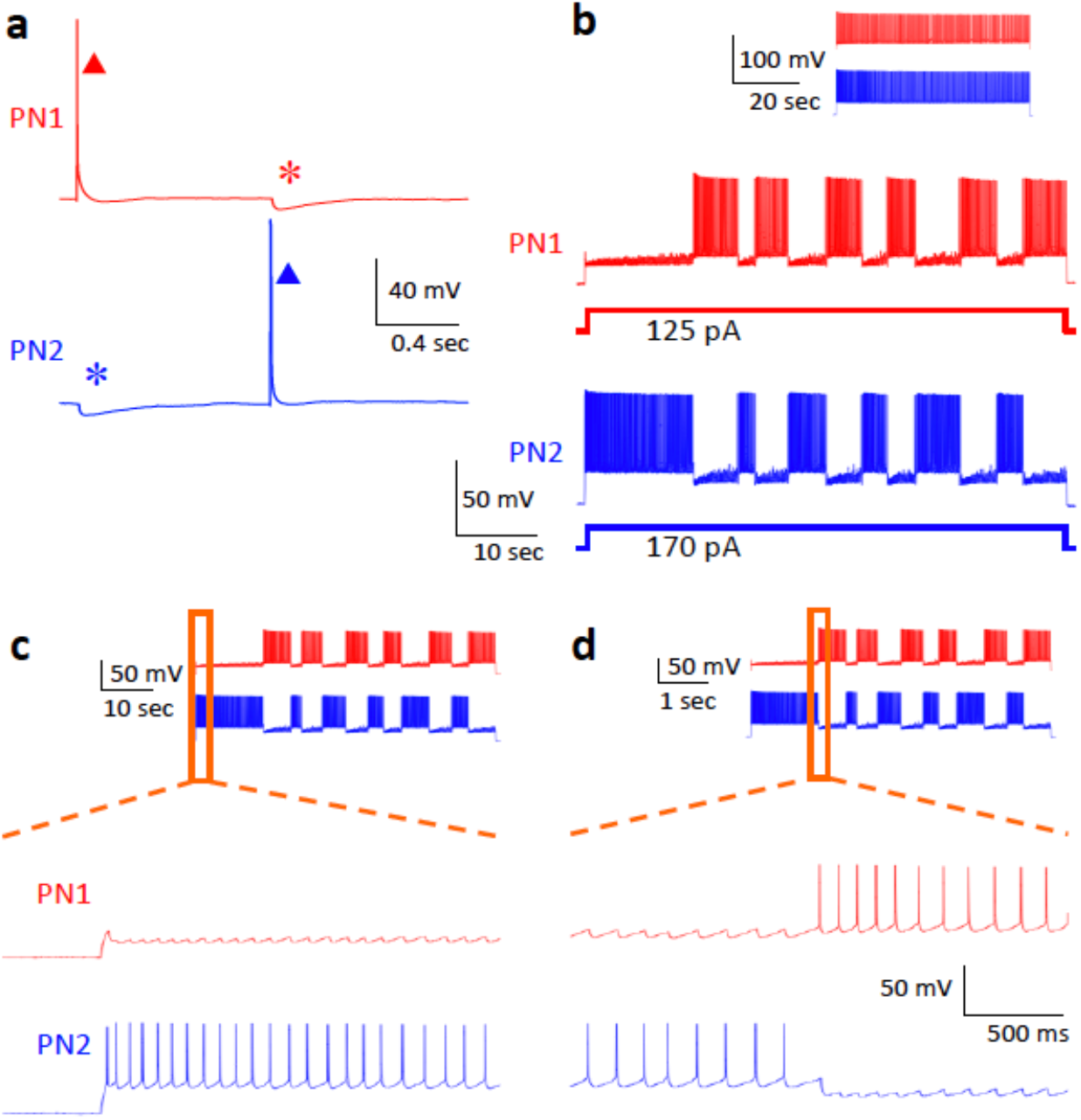
Mutual inhibition between a pair of pyramidal neurons and bi-stable activity. **a**: With the mutual inhibition circuit established, an action potential (red triangle) in the first pyramidal neuron (PN1) triggers an inhibitory postsynaptic potential (IPSP, blue asterisk) in the second pyramidal neuron (PN2); similarly, an action potential of PN2 (blue triangle) causes an IPSP in PN1 (red asterisk). Traces are an average of 5 trials. Baseline membrane potentials were set to −60mV in both neurons. **b**: Continuous injection of depolarization currents into the two pyramidal neurons produces bi-stable activity with alternating dominance between them. MP: membrane potential (mV). MC: membrane current (pA). Inset: The response of the same pyramidal neurons to the same depolarization current injection without the mutual inhibition circuit, showing sustained continuous firing of action potentials. **c**: The part of data (orange rectangle) shown in **b**. Upon the onset of the current injection, both neurons started to depolarize, but fired an action potential first. As a result, PN1 received an IPSP causing PN2 to become dominant and PN1 suppressed. **d**: The part of data (orange triangle) around the time of reversal. The inter-spike interval increased during the dominant period of PN2 due to adaptation. Just after the rightmost action potential of PN2, PN1 got a sufficient time to recover from its IPSP, enabling it to reach its firing threshold before PN2 was able to fire its next action potential. The action potential of PN1 now resulted in an IPSP in PN2 entailing a reversal of dominance.

### Bi-stable activity

When continuous depolarization currents were injected into PN1 and PN2 simultaneously, bi-stable activity with alternating dominance between the two pyramidal neurons was evoked as shown in Fig. 2b. Fig. 2c shows the details of the onset of the response to the current injection on a shorter time scale. Both neurons started to depolarize at the onset but PN2 reached the action potential threshold before PN1 and, hence, PN1 received the evoked IPSP before succeeding to generate an action potential. Thereafter, PN2 showed sustained firing of action potentials and it achieved initial dominance. Note, that an increase of inter-spike intervals in the dominant neuron is visible. Also note that there is a ramp-like slow depolarization of the suppressed neuron (Fig. 2b). The former is a sign of neural adaptation while the latter indicates both the recovery of the neuron from adaptation and the decreasing effect of inhibition from the other neuron. Fig. 2d shows data from when the reversal of dominance occurred. With the continuous increase of inter-spike interval in PN2, PN1 recovered more and more from the received barrage of IPSPs. The inter-spike interval of PN2 eventually became long enough such that the membrane potential of PN1 reached the action potential threshold before PN2 could generate an action potential. Consequently, PN2 received an IPSP evoked by the first action potential of PN1. From then on, PN1 became dominant and PN2 became suppressed.

### Adaptation and dominance durations

To investigate the role of adaptation in the mutual inhibition competition process, we analyzed neurophysiological properties that reflect adaptation: inter-spike intervals and peaks of action potentials are plotted in Fig. 3a, b, respectively, for the example bi-stable activity shown in Fig. 2b. Normalized values are pooled for the “control pairs” (N=93, see Materials and Methods for the definition of the “control pairs”) and plotted over normalized dominance durations in Fig. 3c, d, for inter-spike intervals and action potential peaks, respectively. The results indicate monotonic changes (increase of inter-spike intervals and decrease of action potential peaks over time) while a neuron is dominant. Furthermore, there are clear correlations between the dominance durations and the changes of the inter-spike intervals. We applied a linear regression to inter-spike intervals as a function of time in the dominance duration (see Fig. 9). The slope indicates how quickly the adaptation progresses. As shown in Fig. 3e (for the example shown in Fig. 2b) and Fig. 3f (for the pooled data of the control pairs), the slopes and the dominance durations were inversely correlated (repeated measures ANOVA for the pooled data *F(3,15)=19.518, p<0.0001*). Hence, when adaptation progresses quickly, the dominance duration is bound to be shorter, indicating a role for adaptation in dominance reversals.

**Fig. 3.**
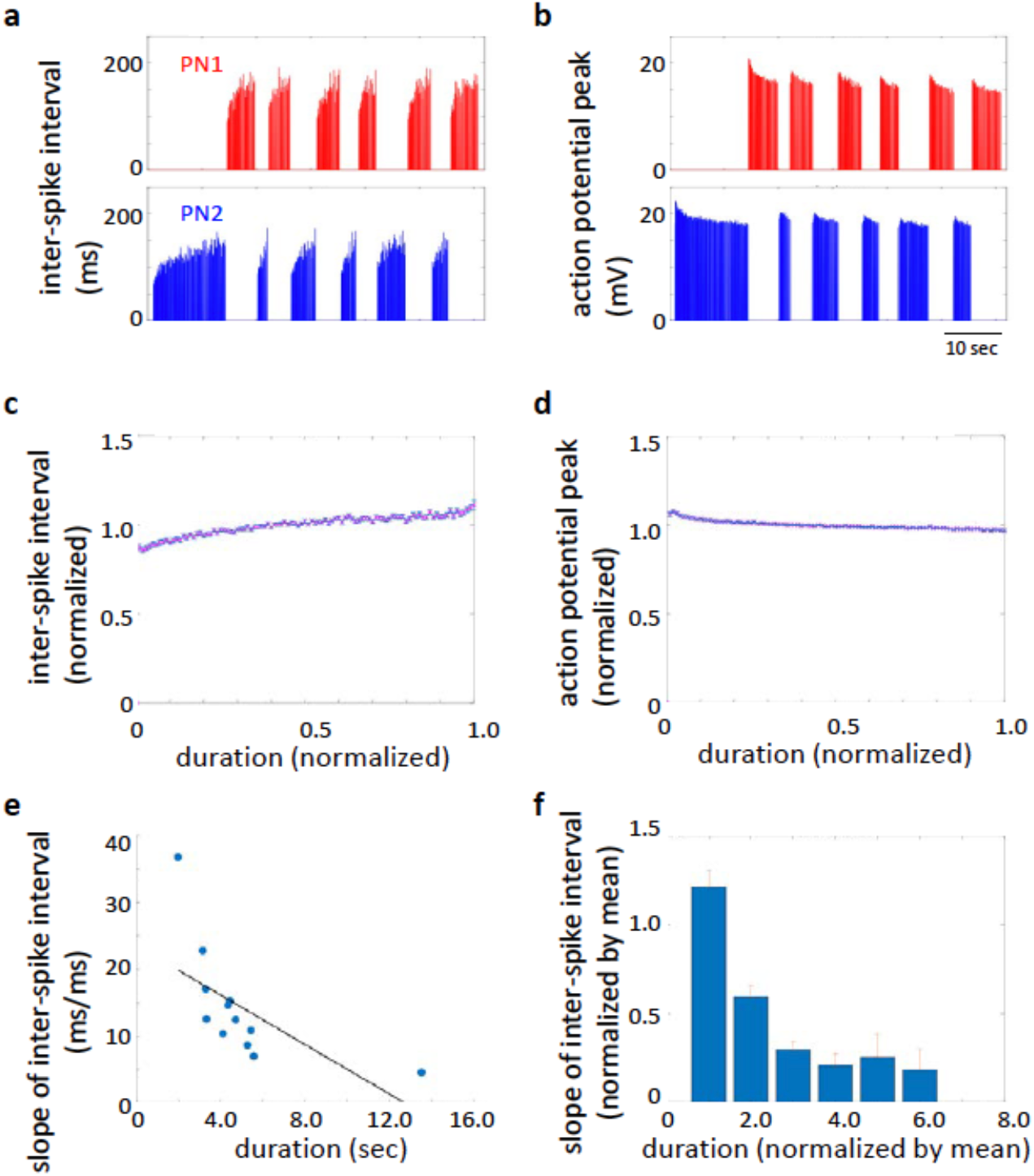
Adaptation of dominant neuron and its correlation to dominance duration. **a-b**: The physiological signatures of adaptation. Inter-spike intervals increase (**a**) and the peaks of action potentials decrease (**b**) due to adaptation during dominance episodes. **c-d**: Average of inter-spike intervals (**c**) and the action potential peaks (**d**) for pooled data of all 93 “control pairs” (see Materials and Methods for the definition). **e**: Slope of inter-spike interval as a function of dominance durations, showing the inverse correlation between them. **f**: Inverse correlation between the slope of inter-spike interval and dominance duration in the pooled data. The dominance durations of individual pairs were normalized by their mean values before pooling. The normalized duration was binned and the pooled data was averaged for the individual bins. Error bars indicate +/− SEM.

### Effect of noise

Because of the stochasticity of dominance durations (Brascamp et al., 2006, p. 2006; Huguet et al., 2014; Kim et al., 2006; Moreno-Bote et al., 2007; Pisarchik et al., 2014) it has been argued that noise plays an important role for the reversal in bi-stable perception. To investigate the role of noise on the dynamics of bi-stability, we implemented an algorithm in the dynamic clamp system to introduce simulated noise of the synaptic conductance (Delgado et al., 2010; Destexhe et al., 2001). The noise was given to both PNs and mINs in the form of random fluctuations of excitatory and inhibitory synaptic conductance (see Materials and Methods for details). Fig. 4a shows the baseline membrane potential of a pyramidal neuron and Fig. 4b shows the result of adding the modeled synaptic noise to it (all at −60mV). Next, the level of noise was changed systematically while the two pyramidal neurons were exhibiting bi-stable activity as shown in Fig. 4c (the parameter sets for different noise level are shown in the table in Fig. 4d). The results indicate that increased noise caused an increase of the reversal rate (*F(19,171)=50.868, p<0.0001*). The pooled data from 15 pairs of pyramidal neurons are shown in Figure 4d.

**Fig. 4.**
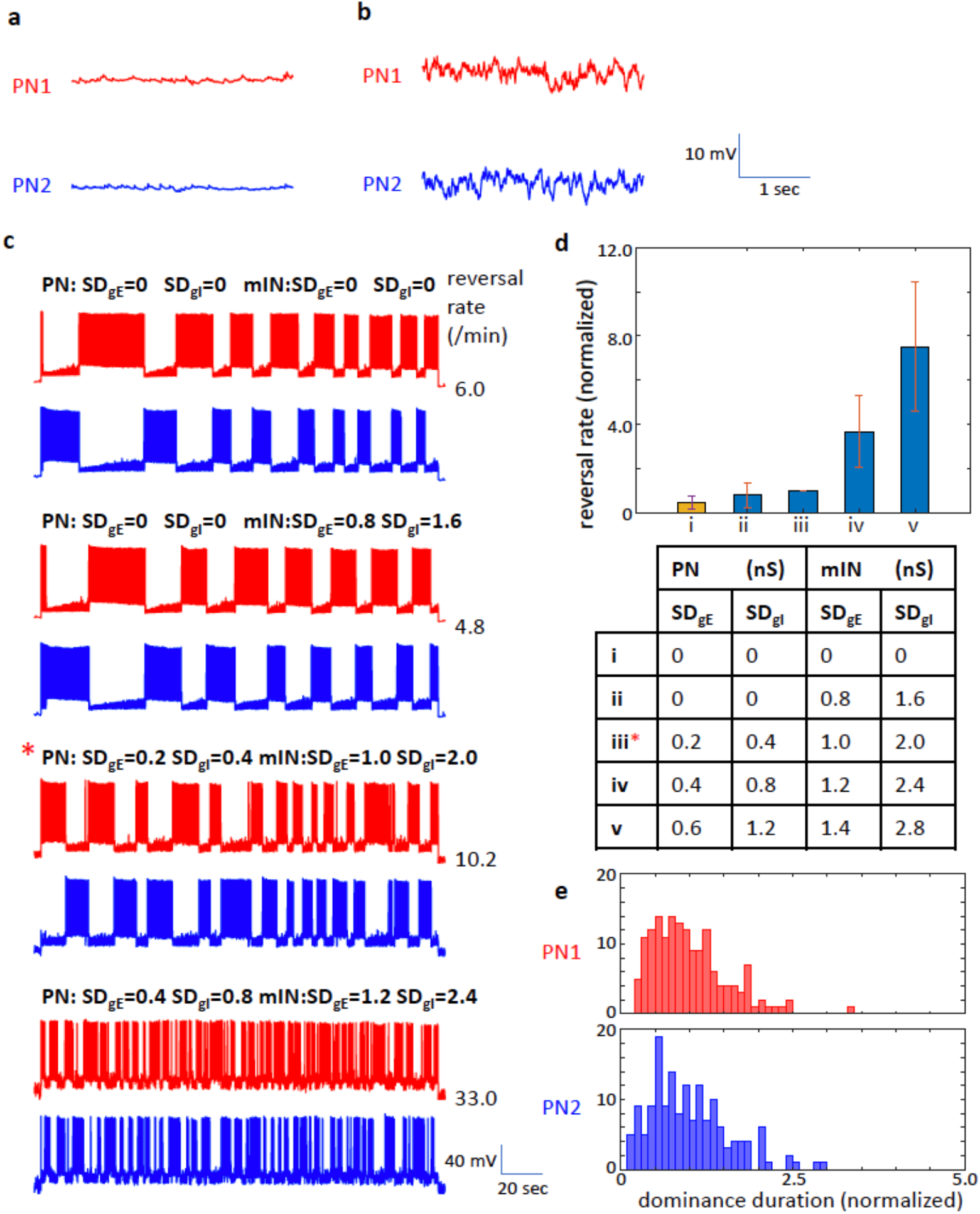
Effect of noise. Model excitatory and inhibitory synaptic noise was applied to the pyramidal neurons and the inhibitory neurons through the dynamic clamp system. **a-b**: Baseline membrane potentials at −60mV without (**a**) and with (**b**) the model noise. **c**: Effect of changing the noise level systematically to bi-stable activity. Increase of the noise resulted in increase of reversal rate (from top to bottom). Noise levels are indicated as standard deviations (SD) of gE and gI (excitatory and inhibitory conductance, respectively, in nS). Asterisk: Data with the “standard” noise parameter set. **d**: Pooled data of the effect of noise (N=15). The reversal rates from the individual pair are normalized by the value at the standard noise parameters (iii) before pooling. Orange bar (i) indicates the data with no model noise. Error bars indicate +/− SEM. The noise parameter sets for i (no model noise), ii, iii (standard noise parameters), iv and v are shown in the table below. The noise level is increased linearly from ii to v. **e**: Histogram of dominance durations for PN1 and PN2 from 10 minutes continuous recording (with the “standard” noise parameters).

It is known that, in brain slice preparations, the amount of synaptic noise in individual neurons is much less than what is observed in intact brain preparations due to the cutoff of axons and lesser spontaneous activity in slice preparations (Destexhe et al., 2001). Therefore, to reproduce the intact brain environment, we use a parameter set of modelled excitatory and inhibitory synaptic noise which will be called the “standard noise parameter set” (asterisk in Fig. 4c and 4d table) from here on. For the rest of the experiments, the standard noise parameter set was used. The histogram of dominance durations of a 600 sec recording of bi-stable activity with the standard parameter set is shown in Fig. 4e. It shows a skewed distribution as stereotypically observed in bi-stable perception. The average of dominance durations and reversal rates of the 15 pairs with the standard noise parameter set were 7.7±5.6sec and 12.0±10.5min^−1^, respectively. These values for the control pairs (N=93) were 8.2±7.8sec and 11.5±10.8min^−1^, respectively.

### Effect of current intensity (“Levelt paradigms”)

A set of widely replicated empirical laws from the perceptual competition literature —known as Levelt’s propositions— describes the relationship between the strengths of two competing stimuli and the dynamics of their bi-stable perception (W. J. M. Levelt, 1965) in terms of dominance, dominance duration, and reversal rate. Furthermore, the paper by Brascamp and Klink (Brascamp et al., 2015) reported a generally accepted updated version of Levelt’s propositions so that the description of bi-stable dynamics covers the full range of stimulus strengths (Levelt’s original propositions were based on the range of stimulation where the stimulus strength of one of the two input signals increased, and hence, the effect of decreasing the strength was not included). To compare the dynamics of the pairs of mutually inhibited pyramidal neurons to the modified version of Levelt’s propositions, we injected sustained depolarization currents and systematically varied (increased and decreased) the strength of the current into one, or both, of the pyramidal neurons (Fig. 5a).

**Fig. 5.**
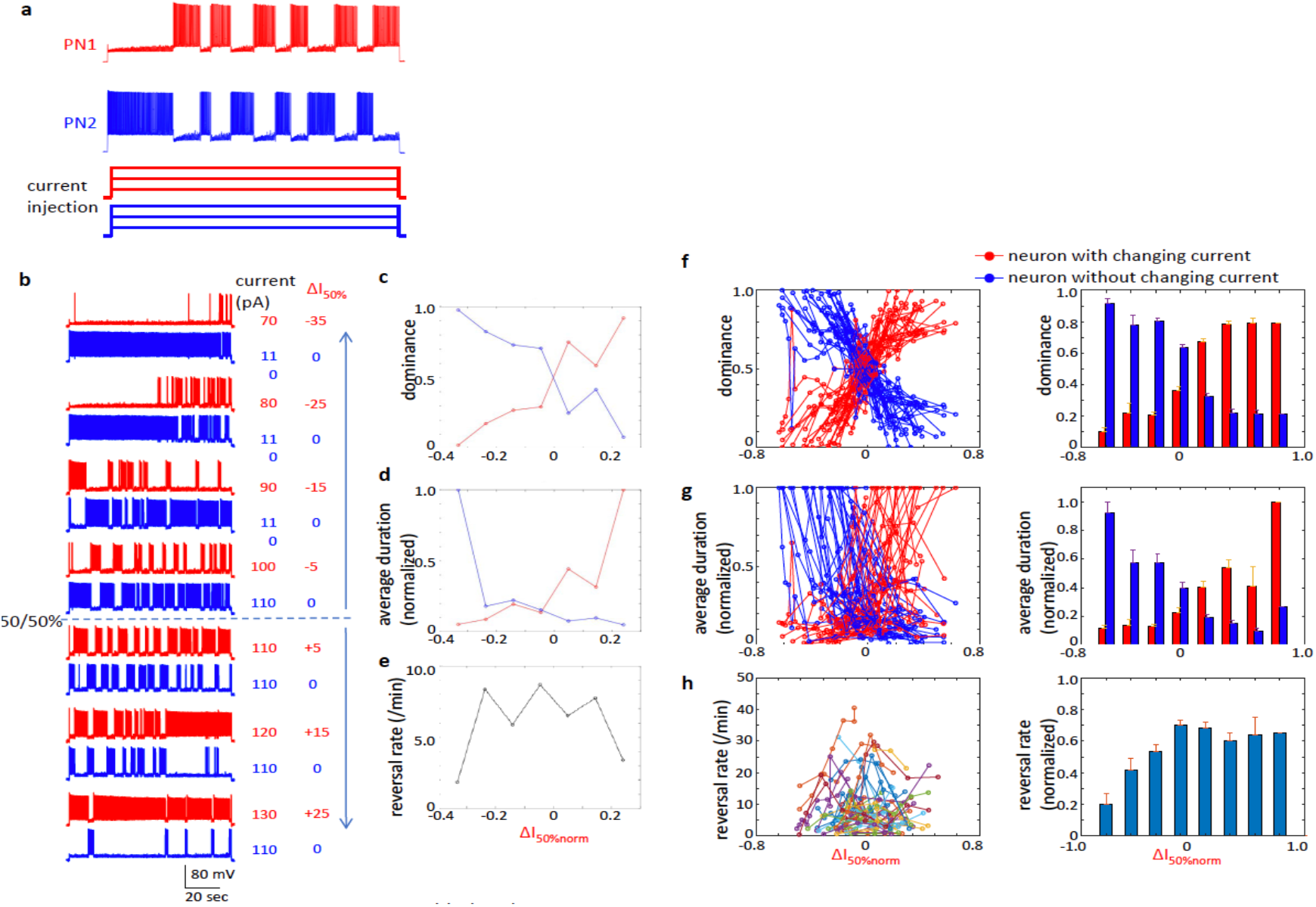
Results of paradigm equivalent to Levelt’s paradigm for proposition I to III. **a**: Schematics of the paradigms equivalent to Levelt’s experimental paradigms for bi-stable perception. The level of injected current to either one or both of the two mutually inhibiting pyramidal neurons was systematically changed (analogous to the change of the contrasts in Levelt’s experiments). **b**: Example data of the experiment equivalent to Levelt’s experimental paradigm for proposition I to III. The level of depolarization current in PN1 was increased (from top to bottom) while the current to PN2 was kept constant. **c-e**: Changes in dominance (**c**), dominance duration (**d**), and reversal rate (**e**) for this pair. PN1 red, PN2 blue. **f-h**: Pooled data (N=46) plotted over the normalized injected current. The dominance durations are normalized for the maximum values of the individual neurons. Red: responses of the neurons that received the changes of the injected current. Blue: responses of the neurons whose injected current was kept constant. Left column: The data of the individual pairs. Right column: The normalized current was binned and the pooled data were averaged for the individual bins. Error bars indicate +/− SEM.

To examine the first three propositions of Levelt, the current injected into one of the two neurons was varied while the current injected into the other neuron was kept constant (Fig. 5b). In total, 46 pairs were recorded with this paradigm. To pool the data, first, the current that would evoke 50% dominance (the total period that one neuron is dominant over the other is equal for both neurons) was estimated (*I*_*50%*_) by linear regression of dominance over the changed current. The change of the current is reported with reference to this control current value (i.e., 0 in abscissa indicates the current pair that would evoke 50% dominance). Hence, in the plots shown in Fig. 5c to 5h, the neurons with the changing injected current is more dominant (“stronger”) on the right side of the plot from 0, while on the left side, they are less dominant (“weaker”).

We first tested Levelt’s proposition I: Increasing stimulus strength for one of the competing stimuli will increase the perceptual dominance of that stimulus. Fig. 5c depicts the change of the dominance ratios of the two pyramidal neurons over injected current (with reference to *I*_*50%*_ of PN1) for the example shown in Fig. 5b. There is a clear trend of increase of dominance of PN1 whose current was increased (red) and of decrease of dominance of PN2 whose current was kept constant (blue). Fig. 5f shows pooled data (N=46) for the dominance ratio, replicating that there is an increasing dominance of the neurons whose currents were increased (red, *F(6,24)=15.558, p<0.0001*), and decreasing dominance for their counterparts whose currents were kept constant (blue, *F(6,24)=15.558, p<0.0001*))xxxcheck thisxxx. This is in line with Levelt’s proposition I.

Levelt’s proposition II states: Increasing the difference in stimulus strength between the two competing stimuli will primarily act to increase the average perceptual dominance duration of the stronger stimulus. Furthermore, the modified Levelt’s proposition II states that the change of stimulus intensity of the non-dominant input is less effective. This means that when the stimulus intensity changes from non-dominant range to dominant range, the effect of the change to average dominant durations is weak in the non-dominant range and strong in the dominant range. In Fig. 5d the change of the average dominance durations is plotted over the changing current for the example shown in Fig. 5b. PN1 shows weak changes of the dominance durations on the left half of the plot where PN1 is weaker than PN2. It shows, however, a steep increase on the right half of the plot where it is stronger than PN2, and vice versa for the other neuron. Hence, in general, the dominant neuron shows a steep increase of the dominance durations with current values deviating further away from *I*_*50%*_. This trend can be seen in Fig. 5g with pooled data for the neurons whose currents were increased (red, *F(6,24)=4.371, p<0.01*)) and for their counter parts whose currents were kept constant (blue, *F(6,24)=7.396, p<0.0001*)xxxcheck againxxx. This is in line with the modified version of Levelt’s proposition II.

According to Levelt’s proposition III: Increasing the difference in stimulus strength between the two competing stimuli will reduce the perceptual alternation rate. Fig. 5e plots the number of reversals for the example shown in Fig. 5b. The pair showed a higher number of reversals for a current close to *I*_*50%*_. Deviating further from *I*_*50%*_ in either direction, the values decreased, in line with Levelt’s proposition III. However, the pooled data (Fig. 5h) show that the response is not symmetric. In fact, some pairs showed an increase of the reversal rate when a neuron is dominant (see Supplement figure, Fig. S1 bottom), in contrast to the example pair of Fig. 5b (and Fig. S1 top). Thus, the pyramidal neuron pairs did not always follow Levelt’s proposition III. Due to the increase in the left half, repeated measures ANOVA indicated a significant effect (*F(6,24)=2.663, p<0.05*).

To examine the fourth proposition of Levelt, the currents injected into both neurons were varied. In total, 32 pairs were recorded with this paradigm. To pool the data, the change of the current is reported with reference to the current that would evoke approximately 10Hz (*I*_*10Hz*_, see Materials and Methods).

Proposition IV states: Increasing stimulus strength of both competing stimuli will generally increase the perceptual alternation rate. In addition, the modified version of proposition IV (Brascamp et al., 2015) noted that this effect may reverse at near-threshold stimulus strengths (i.e. the lower range of stimulation intensity). Fig. 6a shows an example of the effect of increasing the injected currents into both neurons. In Fig. 6b, the number of reversals of this example are plotted over the injected current. Fig. 6c shows pooled data indicating increasing reversal rates (*F(6,30)=4.051, p<0.01*). In addition, there is a small decrease of the reversal rate at the lower range of the stimulation. These results are in line with the modified version of Levelt’s proposition IV.

**Fig. 6.**
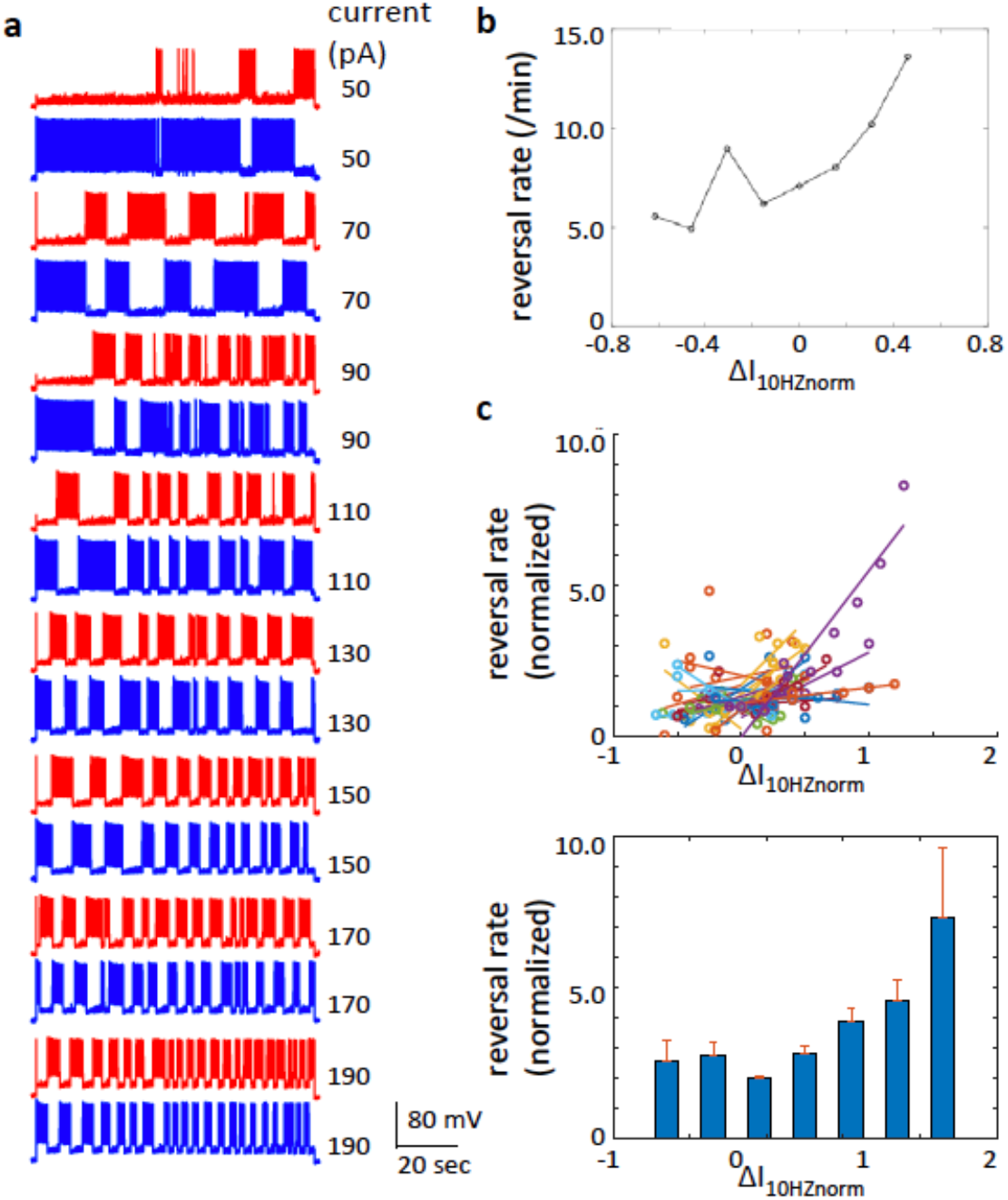
Results of paradigm equivalent to Levelt’s paradigm for proposition IV. **a**: The effect of increasing the depolarization currents simultaneously in both pyramidal neurons (from top to bottom). **b**: The changes of the reversal rate for this pair. **c**: Pooled data of reversal rate (N=32) plotted over the normalized injected currents.

**Fig. 7.**
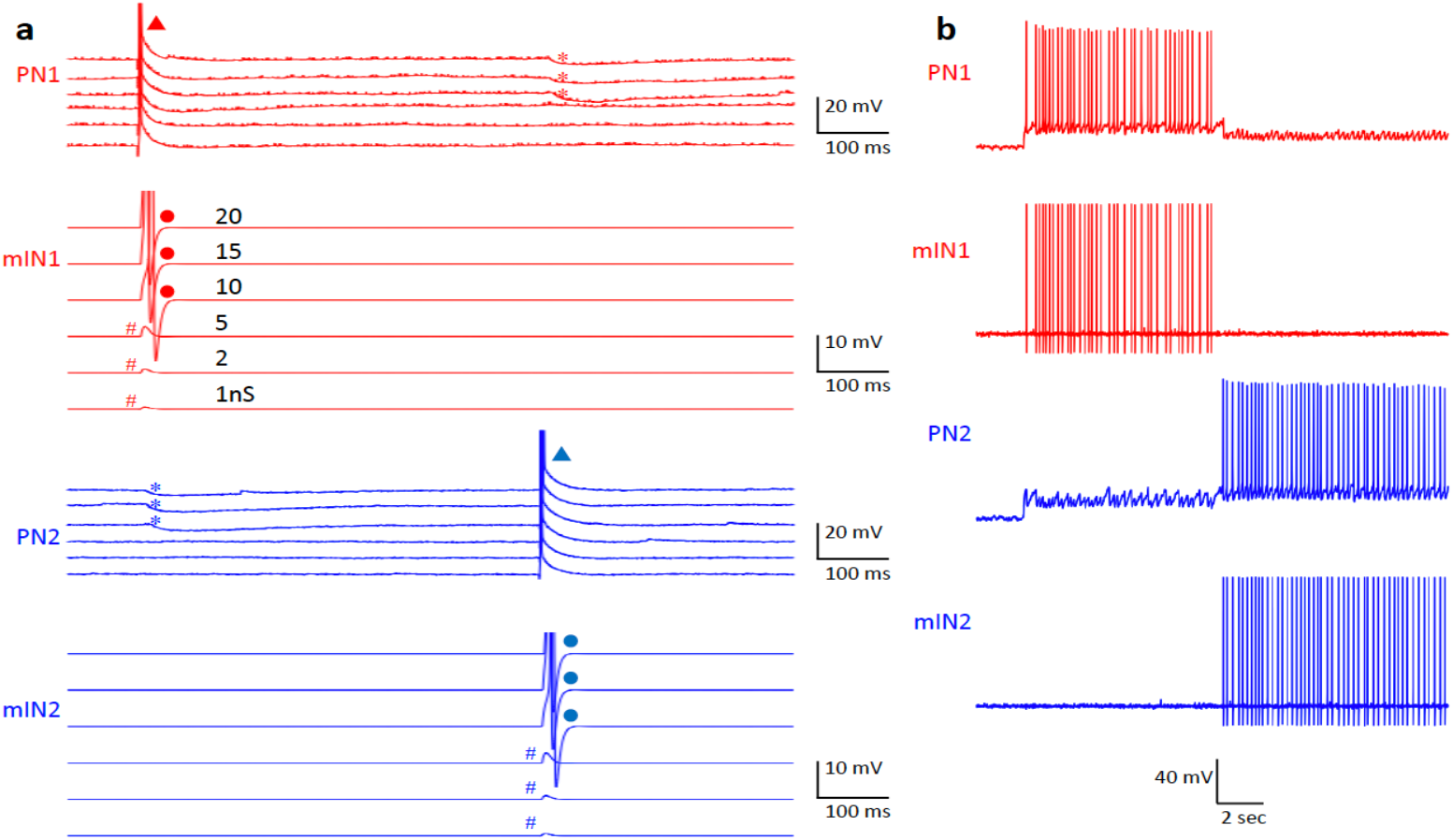
Evoked synaptic events modelled by dynamic clamp. **a**: An action potential in PN1 (red triangle) evoked EPSPs in the partner model inhibitory neuron (mIN1). The synaptic events in mIN1 are shown with six different levels of model synaptic conductance. With the higher synaptic strength, the EPSP evoked an action potential in mIN1 (red disks) causing evoked IPSP in the target pyramidal neuron, PN2 (blue asterisks). When the synaptic strength is in the lower range, it only evoked an EPSP without an action potential in mIN1 (#) and, hence, without an IPSP in PN2. Vice versa from PN2 to mIN2 and PN1. **b**: The activities of PNs and mINs during bi-stable activity.

## Discussion

We developed a system to establish a mutual inhibition connection between two real-life neurons mediated by model neurons and synapses. This system enabled us to evoke bi-stable activity reproducibly in a pair of pyramidal neurons in visual cortex. We analyzed the dynamics of the induced bi-stability, a number of physiological properties, as well as the effects of manipulating the level of background noise and activation level. We compared the dynamics of this pair of neurons with the known dynamics of human bi-stable visual perception. Although our experimental system represents the simplest neural unit of competition and human behavior represents the most complex system, we found that the two systems show striking similarities in their dynamics.

The analyses of the physiological properties during bi-stable activity showed clear signs of neural adaptation of the dominant neurons (Fig. 3). Moreover, we found a link between the variations of inter-spike intervals and dominance durations (Fig. 3e and 3f), indicating a causal link between neural adaptation and the reversals in bi-stable activity. Neural adaptation has only been *assumed* as a key element for bi-stable perception theoretically (Laing & Chow, 2002; Lankheet, 2006; Matsuoka, 1984; Mueller, 1990; Noest et al., 2007; Shpiro et al., 2009; Wilson et al., 2000; Wilson, 1999) or it has been *shown indirectly* in the form of decreased contrast sensitivity (Alais et al., 2010). Our data directly show, in physiological terms, a progression of adaptation during bi-stable activity in pyramidal neurons in visual cortex and its link to the dominance durations.

In addition to neural adaptation, we investigated the effect of neural noise on the dynamics of bi-stable activity. The apparent stochasticity in the sequence of reversals and the skewed distribution of dominance durations (Levelt, 1967) in bi-stable perception led to studies on the role of noise (Baker & Richard, 2019; Brascamp et al., 2006; Huguet et al., 2014; Kim et al., 2006; Moreno-Bote et al., 2007; Pisarchik et al., 2014). To investigate the effect of noise in our experimental model, we incorporated a neuro-computational model of synaptic noise into the dynamic clamp system. In this way, we were able to insert noise into the pyramidal and inhibitory neurons and systematically change the level of noise. We found that an increase of noise caused an increase of reversal rate (Fig. 4). It is known that the synaptic noise found in neurons in brain slice preparations is much less than the noise present in intact brains (Destexhe et al., 2001) or in human brain tissue (Molnár et al., 2008). Hence, we added noise levels equivalent to the noise level in the intact brain (Destexhe et al., 2001). We found that the histogram of dominance durations was right-skewed as is typically found in bi-stable perception.

We showed that when one of the two neurons is dominant, its adaptation progresses and hence the inter-spike interval increases over time. This allows the suppressed neuron to recover from its own adaption and to depolarize more during the ever-increasing inter-spike intervals of the dominant neuron, consequently showing a slowly ramping depolarization. When the depolarized membrane potential comes close to the firing threshold, the noise facilitates the membrane potential to go above the threshold. As a consequence of the action potentials in the previously suppressed neuron, the dominant neuron now receives IPSPs via the disynaptic inhibitory connection and a reversal occurs. Hence, our data elucidate the dynamic interplay between adaptation, noise and mutual inhibition in determining the dynamics of bi-stable activity.

Our experimental model allowed us to separately manipulate the levels of activation of the competing neurons. Hence, it enabled us to compare the effects of changing activation levels in pyramidal neurons to the effects of changes in stimulus strength on the dynamics of bi-stable perception, as originally described in Levelt’s four ‘classic’ propositions (Levelt, 1965). Levelt’s propositions I, II and III make predictions about the changes of dominance, the dominance durations, and the reversal rate, respectively, in response to changes of the stimulation strength in one of the two inputs. Levelt’s proposition IV concerns the change in the reversal rate while the stimulus strengths of both inputs are changed concurrently. The original propositions were modified later (Brascamp et al., 2006, 2015) to cover the whole range of the stimulus strength (dominant and non-dominant ranges). By running paradigms equivalent to these experiments, we found that both systems show striking similarities in their dynamics. Our results strongly suggest that the dynamics reported by Levelt (and its modified version) reflect the dynamics of an underlying neuronal architecture of mutual inhibition circuits.

It is quite intriguing that, although the overall effect of increasing the injections current was the increase of the reversal rate in the paradigm for Levelt’s proposition IV, we observed a small decrease of it in the lower range of the injected currents. As Brascamp and Klink (Brascamp et al., 2015) pointed out, a small deviation of the response from the original proposition by Levelt has been reported by several papers (Curtu et al., 2008; Seely & Chow, 2011; Shpiro et al., 2007). In our experiment, when the injection current was lowered, the firing rate of the neurons became low and the dominant neuron generated action potentials sporadically. As a result, the spike interval became longer, giving room to the suppressed neuron to recover from the inhibition and reach the threshold of action potentials, causing reversal of dominance. On the one hand, in the higher range of injection currents, the reversal occurred because spike intervals of the dominant neuron gradually increased due to adaptation. On the other hand, in the lower range of injection currents, the reversal occurred because of the long spike intervals due to the lower frequency of evoked action potentials. The latter may be potentially a mechanism underlying the small decrease in the lower range of stimulus reported in bi-stable perception.

One exception where our data did not necessarily match the known dynamics of bi-stable perception was the mixed results for the Levelt III paradigm. In this paradigm, some pyramidal neuron pairs showed a decrease of reversal rates when the depolarization current either increased or decreased from the control value, *I50%*, which is in line with Levelt’s proposition III. However, other pairs showed no significant change or an increase of reversal rate when the current was higher than the control (see Supplement Fig. S1 bottom for the examples). Note that the reversal rate is determined by the balance between increased dominance durations of the stronger neuron and decreased dominance durations of the weaker neuron. If the former is more significant, the reversal rate will decrease and if the latter is more significant, it will increase. The mixed results suggest that there may be multiple factors involved in determining the balance. The increase of the firing rate in the stronger neuron with the increase of depolarization current may cause a stronger dominance of the neuron on one hand, and a stronger adaptation of the neuron on the other hand. The latter may prevent the increase of the dominant durations due to the faster decay of the firing rate. Hence, depending on the individually different adaptation properties and the spiking properties of the neurons, the strong activation of the stronger neuron may have caused a decrease of the reversal rate in some cases and an increase in other cases. At systems level, the competition is presumably between populations of neurons rather than single neurons as tested here. Hence, differences in adaptation and spiking properties among the involved neurons may collectively have different impacts on the dynamics of bi-stability. Furthermore, it should be noted that, in the human brain, the input signals go through multiple steps of normalization, starting from the retinae, before reaching the mutual inhibition processes. It may be possible that the activation level of neurons in the human visual system is kept within the range where the fast adaptation occurs in a lesser amount. If this is the case, the strong stimulation would cause the prolongation of dominant durations in the stronger neuron and, hence, cause the decrease of the reversal rate as reported in Levelt III. Therefore, this result may represent an example where emergent properties of bi-stable perception at the behavioral-level differ from the dynamics found in the minimal neural competition unit we investigated.

Regarding the dominance durations, it should be noted that there are short periods when the neuron that has been suppressed fires only one or two action potentials and then becomes suppressed again. Such short events (less than 250 ms) are not considered as a reversal in our analyses, and the dominance durations are determined by neglecting these events (see Fig. 8). Furthermore, there are periods where short events occurred alternatingly between the two neurons with intermingled action potentials from both neurons (Fig. 8). In these periods, none of the two neurons are considered to be dominant. These observations may be linked to known observations in bi-stable perception. It has been reported that human subjects experience short reversal events detected in reflexes (optokinetic nystagmus and pupil dilations) but they are too short to be reported by the subjects (Naber et al., 2011). Furthermore, the intermingled firing of action potentials by the two neurons may be related to the period in bi-stable perception where the perception of the subject is either uncertain or a mixture of the two possible percepts (“composite” or “mixed” perception). The short and the mixture events are potentially important because they may elucidate the neural mechanisms underlying the stochastic properties of bi-stability and decision making processes. This intriguing property of bi-stable neural activity during the transition of the dominances should be investigated further.

**Fig. 8.**
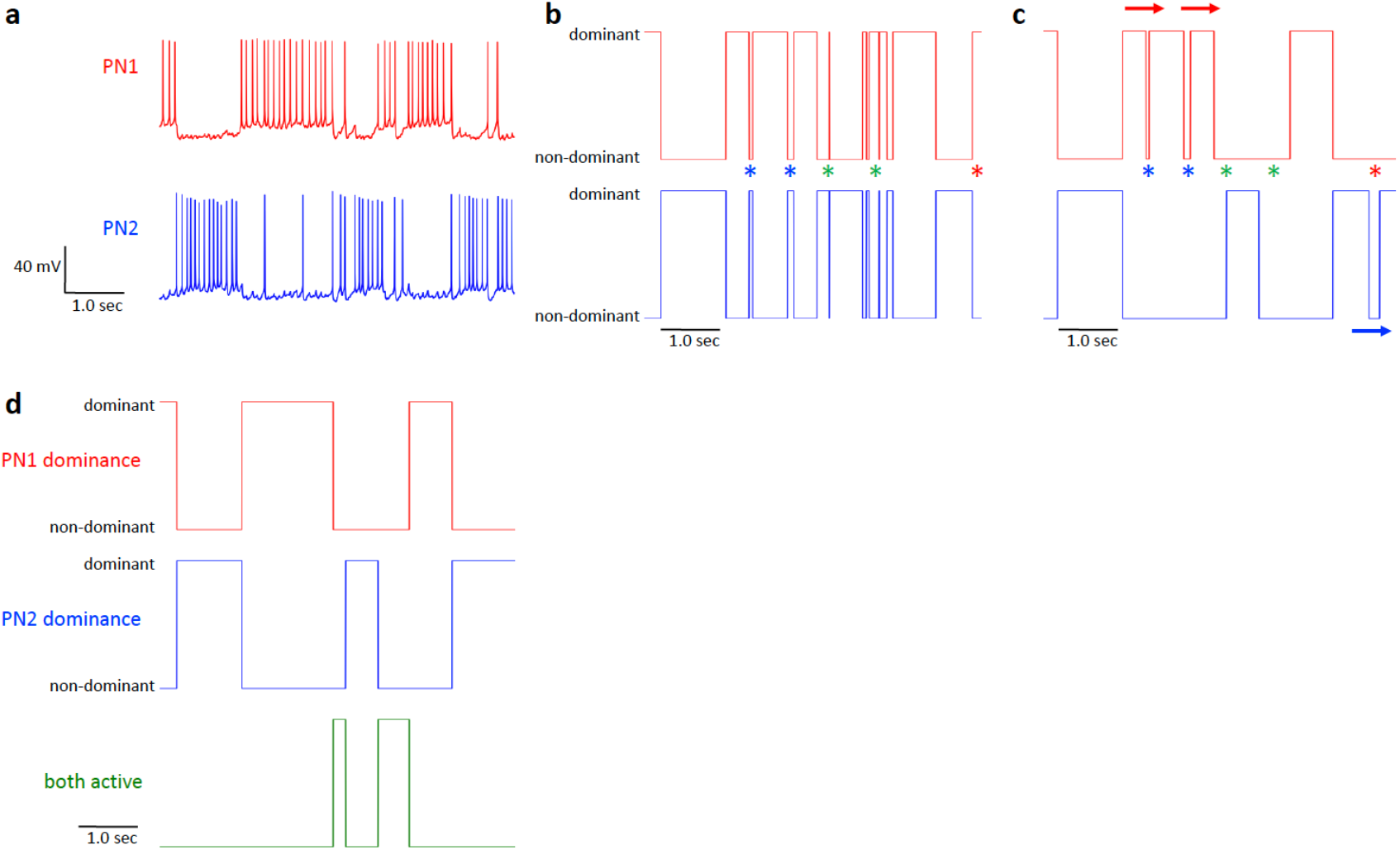
Computation of dominant durations. **a**: A part of a recording of bi-stable activity. **b**: First step computation of dominance durations. Here, continuous firing of action potentials in one neuron until an action potential occurs in the other neuron is considered as a tentative dominance duration of the first neuron. Hence, the dominant durations of the two neurons are mutually exclusive. Note that there are short dominant durations (blue asterisks for PN2 and red asterisk for PN1). There are also series of alternations of short dominant durations between the two neurons (green asterisks). **c**: Dominance durations after choosing only long durations (longer than 250 ms). This process results in short lags between the dominance durations (blue and red asterisks). There are also the intervals that are not assigned to either of the neurons corresponding to the period marked with green asterisks in **b**. The short lags are not considered as reversals and, hence, the previous dominance is considered to continue (arrows). **d**: These processes result in the final dominance durations without short durations. And the periods not assigned to neither of the neurons are assigned as “both active” (bottom).

**Fig. 9.**
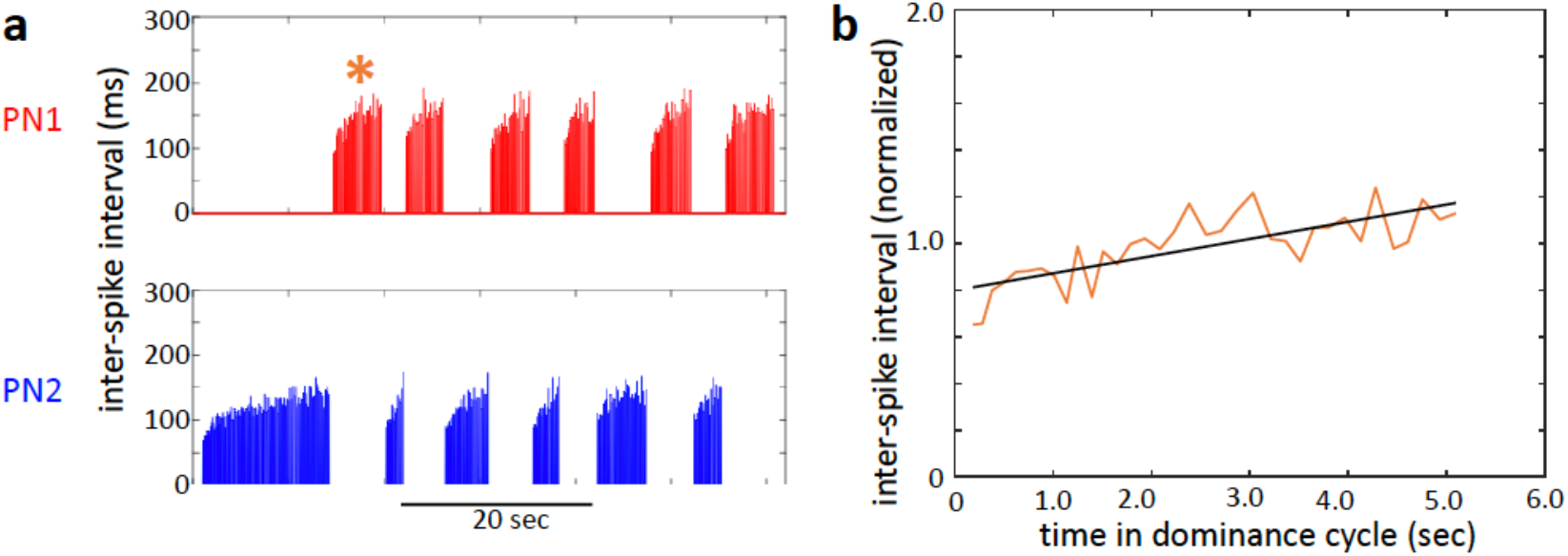
Linear fitting to progress of inter-spike interval. **a**: Inter-spike intervals of an example shown in Fig. 2B. **b**: Linear regression (black) of inter-spike intervals (orange plot) taken from the first cycle of PN1 (orange asterisk in **a**) plotted over time from the onset of its dominance duration.

Concluding, our experimental model provides a platform for investigating the dynamics of a theoretically derived neural circuit in real-life neurons. Our data showed that even the simplest neural competition circuit, between two individual pyramidal neurons, already reproduces many aspects of dynamics of bi-stable perception in human perception. Our study using the novel approach reported here provides a platform to investigate further how elementary neural competition units are integrated to execute system-level bi-stable dynamics.

## Materials and Methods

Experiments were performed at the Brain Science Institute (Tamagawa University, Japan), and the Donders Institute for Brain, Cognition and Behavior, (Radboud University, The Netherlands). The experimental animal procedures were approved by the Animal Research Ethics Committee of Tamagawa University (animal experiment protocol H29/08) and the Animal Ethics Committee of the Radboud University Nijmegen, under DEC application number 2018-0016 (Nijmegen, the Netherlands). The procedures are in accordance with the Guidelines for Animal Experimentation in Neuroscience (Japan Neuroscience Society) and the Dutch legislation.

### Brain slice preparation

Brain slices were prepared from the occipital part of the mouse brain that includes the visual cortex (strain C57BI6/J, age p12 to p24). Mice were anesthetized deeply using isoflurane in an induction chamber. Following deep anesthesia, mice were quickly decapitated and the brain was removed from the skull in a small container with chilled “cutting solution”. For this process, the solution of either one of the following compositions was used (in mM): 125 NaCl, 25 NaHCO_3_, 2.5 KCl, 1.25 NaH2PO4, 1 CaCl_2_, 2 MgCl_2_, 25 D-glucose, or 75 sucrose, 87 NaCl, 25 NaHCO_3_, 2.5 KCl, 1.25 NaH_2_PO_4_, 0.5 CaCl_2_, 7 MgCl_2_, 25 D-glucose, both saturated with 95% O_2_, 5% CO_2_. Then, the brain tissue was glued on to the cutting stage of a vibratome (VT1000S, Leica, Germany, or Microm HM 650V, Thermo Scientific, USA), submerged in the cutting solution above. Coronal or angled-coronal (Dong et al., 2004) sections of 300~400μm thickness were cut and stored in an incubation chamber in 32~34°C for at least 30 min, and then stored at room temperature until use.

### Double whole-cell recordings

Slices were transferred to a recording chamber on a microscope stage and were superfused with artificial cerebrospinal fluid, ACSF, maintained at a constant temperature (32~34°C). ACSF had the following composition (in mM): 125 NaCl, 25 NaHCO_3_, 2.5 KCl, 1.25 NaH_2_PO_4_, 2 CaCl_2_, 1 MgCl_2_, 25 D-glucose, saturated with 95% O_2_, 5% CO_2_. The location of V1 was identified under the microscope (Olympus, Japan) equipped with DIC-IR (differential interference contrast – infrared). Layers of visual cortex were identified and the point where layer 5 starts thickening, going from medial to lateral, was used as a landmark of the border between V1 and LM (lateromedial area, Wang and Burkhalter, 2007, equivalent to V2, Fig. 1c). All recordings were made from the region medial from the landmark. Under high magnification with x40 objective, pyramidal neurons in layer 2/3 were identified by their stereotypical morphology. In some cases, the recorded neurons were filled with biocytin and post-experimental process indicated that, in all cases, they were pyramidal neurons in layer 2/3 (see below). Two neurons separated by at least 150μm distance were selected to reduce the probability of choosing connected pairs. Furthermore, experimental protocols were performed to check for monosynaptic (paired-pulse injection at 10Hz to one of the neurons to evoke action potentials) and disynaptic connections (Kapfer et al., 2007; Silberberg & Markram, 2007) (100Hz 11 pulses injection to one of the neurons to evoke a train of action potentials). None of the pairs reported in this paper were connected.

Pipettes for patch clamp recordings were pulled from borosilicate thin glass capillaries (TW150-4, WPI, USA) and filled with a filtered intracellular solution with the following composition (mM). 130 K-gluconate, 10 KCl, 4 ATP-Mg, 0.3 Na-GTP, 10 HEPES, 10 phosphocreatine. For phosphocreatine, either 10mM Na_2_-phosphocreatine or a mixture of 5mM Na_2_-phosphocreatine and 5mM tris-phosphocreatine was used. The osmolarity of the solution was adjusted to 290~300Osm by either Osmotron-5 (Orion Riken Co., Japan) or Semi-Micro Osmometer K-7400 (Knauer, Germany) and the pH was adjusted to 7.2. The final resistance of the pipettes was 7~9MΩ. In some cases, biocytin was added to the pipette solution (2.5~5mg/ml) to visualize the recorded neurons post-experimentally. Recordings were carried out using either two Axopatch 200B amplifiers or a Multiclamp 700 amplifier (both Molecular Devices, Sunnyvale, CA, USA). Data were lowpass filtered at 10kHz and were digitized at 20 kHz using a Digidata A/D board model 1440A. Data were captured using the Clampex program suite (Molecular Devices, USA). Series resistances were constantly monitored by injecting a −100pA pulse in current-clamp configuration. Series resistances were balanced via a bridge circuit.

### Cell identification

To visualize the pyramidal neuron pairs that were recorded, they were filled with biocytin by diffusion (N=9). After the recording (approximately 30 to 60 minutes), the slices were kept in 4% paraformaldehyde in phosphate buffer solution, PBS, (0.1 M, pH 7.2) and were kept at 4°C. After washing the tissue with PBS, it was quenched with 1% H_2_O_2_ in 10% methanol and 90 % PBS for 5 minutes. The tissue was washed with PBS and permeabilized with 2% Triton X-100 in PBS for 1 hour and then put in ABC solution (ABC Elite Kit, Vector, USA) overnight at 4°C. After washing the tissue with PBS and then with Tris buffer (0.05M), it was processed with DAB solution (0.5g/l in 0.05M Tris buffer) and 1%H_2_O_2_ was added to enhance the reaction. After verifying the visualization of neurons, the tissue was washed by PBS and then mounted to glass slides with a mounting medium (Aquamount, Vector, USA).

### Dynamic clamp

A modified version of the dynamic clamp system StdpC (spike timing dependent plasticity clamp)(Nowotny et al., 2006) was used to establish the connections between recorded neurons and model neurons with model synapses. The communication between the amplifier and StdpC was mediated by a National Instruments A/D board, model PCIe-6321. Dynamic clamp is a method whereby a modelled conductance, e.g. a synaptic or ionic conductance, is computed based on the measured membrane potential of a neuron, then injected into that neuron in real time with a patch clamp electrode. Unlike other dynamic clamp systems which operate at fixed frequencies, StdpC does not require a real-time operating system, relying instead on precise measurement of the time elapsed in each measure-compute-inject cycle to perform the numerical integration of its models.

Besides numerous improvements to the software interface, the following major additions were made to the previous version of StdpC (Nowotny et al., 2006). A passive membrane model was added, which can be augmented with models of ionic and synaptic conductances to form completely synthetic neuron models. To stabilize numerical integration of such models at StdpC’s unpredictable and varying sampling frequency, the clamp cycle was upgraded from explicit Euler to a Runge-Kutta integration scheme of order 4/5. A number of performance enhancements were made to ensure high-frequency, and thus high-fidelity, updates to the injected current. A delay mechanism was added to the synapse models, allowing the simulation of conduction and synaptic delays. Finally, a model of synaptic background noise was added, reproducing the synaptic bombardment we would expect to see in vivo with statistically equivalent, randomly generated inhibitory and excitatory currents, as described in the section on noise below. The upgraded version of StdpC (StdpC version 6.1) is available at github.com (github.com/CompEphys-team/stdpc, DOI 10.5281/zenodo.3492203).

A custom-made summing circuit was used to combine the command signal from StdpC and the one from Clampex software, and the combined command signal was fed to the amplifier.

Hodgkin-Huxley models of ionic channels (conventional sodium, delayed rectifier potassium, and Kv3 potassium channels) were given to the model inhibitory neuron (membrane capacitance 0.2115nF, leak conductance 63.462nS, equilibrium potential for the leak conductance −70mV (Pospischil et al., 2008)). A Kv3 channel was added to simulate fast spiking inhibitory neurons(Lien & Jonas, 2003). The models are based on an “α/β formalism” as follows (see github.com/CompEphys-team/stdpc/tree/master/manual).

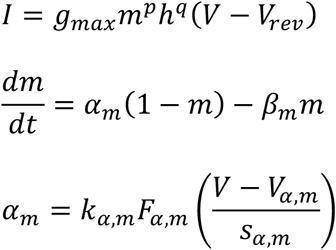

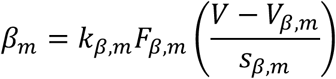

(and analogous for *h*).

Here, *m* and *h* are activation and inactivation variables. *g*_*max*_ is the maximum conductance of the ion channel and *V*_*rev*_ is the reversal potential of the ion. The form of the function *F* is either one of the three below.

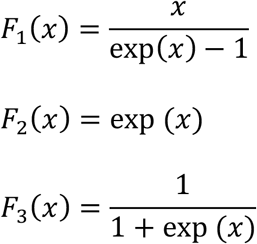

For the potassium channels, the formalisms are the same, except that no inactivation components are included. The form of the function *F* and the parameters for *α* and *β* for the individual components are as summarized in Table 1 (top). These parameter values were taken from Pospischil *et al.* (Pospischil et al., 2008) for basic membrane properties, from Hodgkin & Huxley(Hodgkin & Huxley, 1952) for sodium and delayed rectifier potassium channels and from Lien & Jonas (Lien & Jonas, 2003, p. 3) for KV3 channel.

**Table 1.**
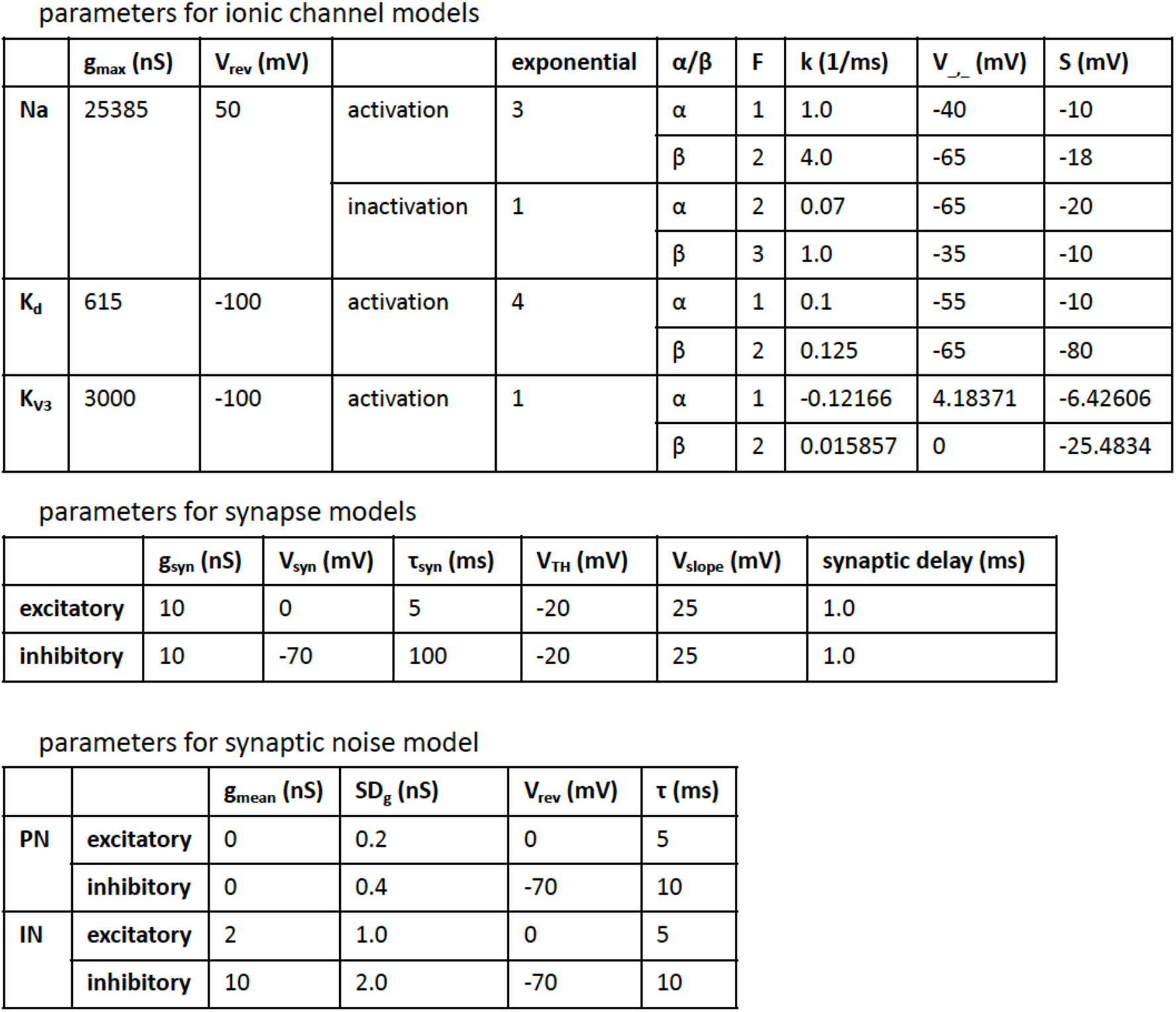
Parameter sets for neuron models. Parameter sets for modelled ionic channels (top), synaptic conductance (middle) and synaptic noise (bottom).

| | $g_{max}$ (nS) | $V_{rev}$ (mV) | | exponential | $\alpha/\beta$ | F | k (1/ms) | $V_{\infty}$ (mV) | S (mV) |
| --- | --- | --- | --- | --- | --- | --- | --- | --- | --- |
| Na | 25385 | 50 | activation | 3 | $\alpha$ | 1 | 1.0 | -40 | -10 |
| | | | | | $\beta$ | 2 | 4.0 | -65 | -18 |
| | | | inactivation | 1 | $\alpha$ | 2 | 0.07 | -65 | -20 |
| | | | | | $\beta$ | 3 | 1.0 | -35 | -10 |
| $K_d$ | 615 | -100 | activation | 4 | $\alpha$ | 1 | 0.1 | -55 | -10 |
| | | | | | $\beta$ | 2 | 0.125 | -65 | -80 |
| $K_{v3}$ | 3000 | -100 | activation | 1 | $\alpha$ | 1 | -0.12166 | 4.18371 | -6.42606 |
| | | | | | $\beta$ | 2 | 0.015857 | 0 | -25.4834 |

parameters for synapse models
| | $g_{syn}$ (nS) | $V_{syn}$ (mV) | $\tau_{syn}$ (ms) | $V_{TH}$ (mV) | $V_{slope}$ (mV) | synaptic delay (ms) |
| --- | --- | --- | --- | --- | --- | --- |
| excitatory | 10 | 0 | 5 | -20 | 25 | 1.0 |
| inhibitory | 10 | -70 | 100 | -20 | 25 | 1.0 |

**Table 1** Parameter sets for neuron models
| | | $g_{mean}$ (nS) | $SD_g$ (nS) | $V_{rev}$ (mV) | $\tau$ (ms) |
| --- | --- | --- | --- | --- | --- |
| PN | excitatory | 0 | 0.2 | 0 | 5 |
|  | inhibitory | 0 | 0.4 | -70 | 10 |
| IN | excitatory | 2 | 1.0 | 0 | 5 |
|  | inhibitory | 10 | 2.0 | -70 | 10 |

Conductance of excitatory and inhibitory synaptic events were modeled using the ChemSyn model in StdpC, following the equations and parameters described below.

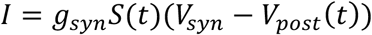

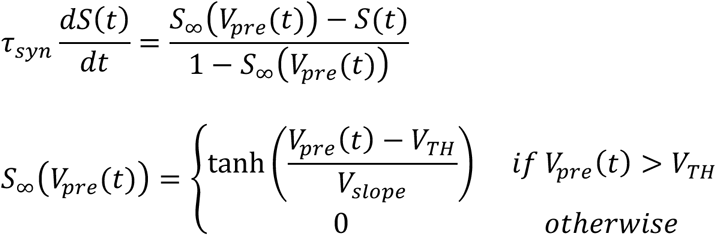

Parameters for excitatory and inhibitory synapses are shown in Table 1 (middle). *G_syn* for EPSP was selected so that it evokes an action potential in mINs (Fig. 7), and *g_syn* and *τ_syn* for IPSP were selected to ensure strong enough suppression of target PN. The synaptic delay was set to 1ms in all cases, and no synaptic plasticity was included in the model.

### Disynaptic mutual inhibition connections

Establishment of a mutual inhibition circuit was verified as follows. Injection of a brief (1ms) depolarization current (1500~2000pA) to one of the pairs of pyramidal neurons evoked an action potential (red and blue triangles in Fig. 7a), which triggered an excitatory synaptic conductance in the model inhibitory neuron. This synaptic event evoked an EPSP in the inhibitory neuron. As shown in Fig. 7a, when *gMax* was set to 10nS or higher, the EPSP evoked an action potential (red and blue disks). This action potential in the inhibitory neuron triggered an inhibitory synaptic conductance, which was fed to the postsynaptic pyramidal neuron as an injected IPSC via the amplifier, giving rise to a corresponding IPSP (blue and red asterisks). Fig. 7a shows that an action potential was first evoked in the pyramidal neuron 1 (PN1) and the pyramidal neuron 2 (PN2) received an IPSP. Later, an action potential was evoked in PN2 that resulted in an IPSP given to PN1, illustrating that the mutual inhibition circuit was established between the two pyramidal neurons by this system. As shown in Fig. 7b, the inhibitory neurons show trains of action potentials corresponding to the action potentials of presynaptic pyramidal neurons during bi-stable activity.

### Bi-stable activity

Bi-stable activity is evoked by the following protocol. First, before the dynamic clamp mediated model circuit is switched on, depolarization currents that evoke action potentials at approximately 10 Hz in the two neurons are determined separately. Next, the model circuit is switched on to activate the mutual inhibitory connection, and the depolarization currents as determined above are injected. In most cases, this already produces bi-stable activity in the pair (unless one of the neurons is 100% dominant). However, every neuron has different firing patterns, different degrees of responses to given synaptic inputs, and different sizes of action potentials (which influence the strength of postsynaptic events). As a result, the bi-stable activity often does not show equal dominance between the two neurons even though the firing rates are equivalent between them. Therefore, in the case that it is necessary to find the current pair where the dominance of the two neurons are approximately equal (50% dominance point), the currents are further adjusted by either increasing the current in the weaker neuron or decreasing the current in the stronger neuron.

Dominance, dominance durations, and reversal rates were calculated using custom Matlab (MathWorks, USA) scripts. Unlike behavioral studies, in which a dominant percept is indicated as a continuous signal (by button press), the dominance of a neuron is signaled by sustained repetitive firing of action potentials. Hence, we defined the “dominance duration” of a neuron as follows (illustrated in Fig. 8). First, a continuous firing of action potentials in one neuron until an action potential occurs in the other neuron is considered as a tentative dominance duration of the neuron (Fig. 8b). Hence, at this stage, the dominance durations of the two neurons are mutually exclusive. Note that there are short dominance durations (blue asterisks for PN2 and red asterisk for PN1). There are also a series of alternations of short dominance durations between the two neurons (green asterisks). Next, dominance durations shorter than 250ms are eliminated (Fig. 8c). This process results in short lags between the dominance durations (blue and red asterisks). The occurrence of the short lag is not considered as reversal and, hence, the previous dominance is considered to continue (arrows). These processes result in the final dominance durations without short durations (Fig. 8d). Note that there are also the intervals that are not assigned to either of the neurons corresponding to the period marked with green asterisks in Fig. 8c. This is because alternating short durations occur between the two neurons during these periods (Fig. 8c green asterisks). These periods are assigned as “both active” (Fig. 8d bottom). Dominance and reversal rates were computed based on this definition of dominance durations. “Dominance” of a neuron is defined as the ratio of total dominance durations of the neuron (sum of all dominance durations of the neuron) divided by the sum of the total dominance durations of both neurons. A reversal is defined as the dominance switching from one neuron to the other, regardless of the presence or absence of a “both active” phase during the switch.

Special attention was paid to the recording conditions. If the following criterion were not met, the recording was halted: The overshoot of action potential should be higher than 10mV, and changes in the size of the action potential, in series resistance, and in firing rate to a given depolarization current should be less than 15% during data collection.

### Analysis of adaptation

Inter-spike intervals and the peaks of action potentials were estimated with custom Matlab scripts. Upon detection of action potentials inter-spike intervals and the peaks were measured. These values were plotted against time to visualize the progress of adaptation within individual dominance episodes (Fig. 3a, b). To pool the data, first, the time from the onset of the dominance cycle to the end of this cycle was normalized by dividing it by the cycle’s dominance duration (for the individual cycles of the individual pairs), resulting in the normalized time ranging from 0 to 1. Second, inter-spike intervals and the magnitude of action potential peaks were normalized by the first values of the individual cycles. Third, the normalized values across all pairs were sorted into bins of size 0.01. Finally, the mean and standard deviation of all inter-spike intervals and action potential peaks in a given bin were plotted against the normalized time (Fig. 3c and 3d). As an indicator of the progress of adaptation, inter-spike intervals (normalized by the mean of individual pair) was plotted over time from the onset of each dominance cycle and linear regression was applied to the plot (Fig. 9). This resulted in slope values that indicated the change of inter-spike intervals. To pool the data, the dominance durations of individual pairs were normalized by their mean values and the slopes, normalized by the mean values of individual neurons, were plotted over the normalized duration (Fig. 3f).

### Effect of noise

To investigate the effect of noise on the dynamics of bi-stable activity, synaptic background activity was simulated according to the model by Destexhe *et al.* (Destexhe et al., 2001). In their simulation, random walk-like fluctuations of membrane conductance were modeled by applying the Ornstein-Uhlenbeck model of Brownian motion (Uhlenbeck & Ornstein, 1930). Their formalism of synaptic noise was implemented in the StdpC dynamic clamp system. The evolution of the simulated synaptic noise depends on the noise time constant *τ*, which controls noise color, as well as the mean *gmean* and standard deviation *SDg* of the noise, and is modeled as follows:

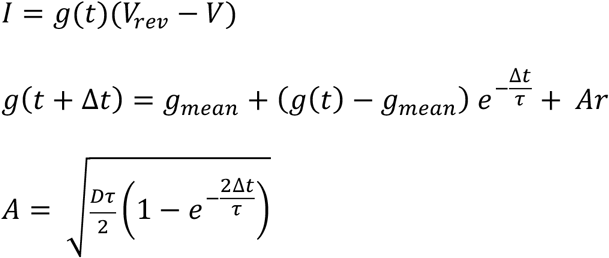

Here, *r* is a pseudo-random number drawn from a normal distribution with mean 0 and standard deviation 1, and the noise diffusion coefficient *D* is related to the noise standard deviation as follows:

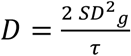

Excitatory and inhibitory synaptic noise are modeled separately. The level of noise is expressed as the standard deviation *SDg* of the synaptic conductance and systematically manipulated, whereas the average conductance *g*_*mean*_, which functions as a constant current offset, remained unchanged. The amount of noise given to mINs was larger than that given to PNs because PNs already have intrinsic synaptic noise (Fig. 4a) from their presynaptic neurons within the brain slice. The standard parameter set (used as default unless mentioned otherwise) for the noise is shown in Table 1 (bottom).

In the experiments for the effect of noise level and the effect of activation level (below), the length of each trial was 200 sec with 193.5 sec long depolarization current.

### Paradigms equivalent to Levelt’s experiments

For our experiments associated with the classic behavioral experiments of Levelt (Levelt, 1965), we systematically varied the strength of the sustained depolarization current into one, or both, of the pyramidal neurons. Concerning Levelt’s proposition I to III, only one of the two currents (randomly selected) was altered. The change of the current was made by steps of 10 or 20pA.

In the analyses, the current that would evoke 50% dominance, called *I*_*50%*_, was estimated by linear regression of dominance over the changing current. The change of the current is reported with reference to this control current value, defined as follows.

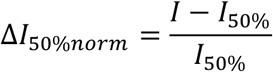

Hence, in the plots in Fig. 5c to 5h, the right side from x=0 indicates that the neuron that received the changing current was more dominant (stronger) than the other neuron and the left side indicates the former being weaker than the latter. Before pooling the data (N=46) for average durations, average dominance durations of individual trials were computed and were divided by the maximum average duration within individual neuron. To pool the data for the reversal rate, data were normalized by the maximum reversal rate of the individual pair (Fig. 5h). Concerning Levelt’s proposition IV, both currents were modified. First, a current pair that evoked a 10Hz firing rate in the two neurons was found. If necessary, the current was adjusted until the current pair evoked approximately 50% dominance. This current pair was considered as a control and is called *I*_*10Hz*_(it is called as such for convenience although the current pair did not always evoke 10Hz firing). Next, in one of the two neurons, the current was changed with 10 or 20pA steps and the current for the other neuron was changed proportionally. To pool the data, the change of the current is reported with reference to *I*_*10Hz*_, defined as follows.

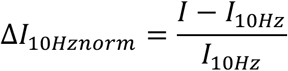

To pool the data for the reversal rate (N=32), data were normalized by the reversal rate of the individual pair when the control current pair was used (Fig. 6c).

To make the bar plots of the pooled data (Fig. 5f to h, right, Fig. 6c bottom), the Δ*I* was binned and the values in the individual bins were averaged. The order of trials with different current pairs was pseudo-randomly chosen.

For statistical analysis, repeated measures analysis of variance (ANOVA) was applied using SPSS Statistics (IBM, USA). Pairs with the standard noise parameter set for the experiment of noise (N=15), pairs with injected current of *I*_*50%*_ in Levelt I to III paradigms (N=46), and pairs with injected current of *I*_*10Hz*_ in Levelt IV paradigm (N=32) are collectively called a “control pair” and statistical analyses were performed on these 93 pairs to report basic properties of bi-stability and adaptation. Error bars in the plots are +/− SEM.

All data and Matlab codes for data analyses are published at Radboud University data repository site with URL as below. https://data.donders.ru.nl/collections/di/dcn/DAC_626810_0008_424?93

## Acknowledgements

We would like to thank Dr. Ginamaria Maccaferri (Northwestern University, Chicago, USA) for supporting the project during the time of piloting, Dr. Andreas Burkhalter (Washington University, St. Louis, USA) for providing detailed information of the anatomy of mouse visual cortex, Dr. Yoshikazu Isomura (Tamagawa University, Machida, Japan) for supporting to conduct the project and the anatomical analysis at Brain Science Institute, Dr. Nael Nadif Kasri and Dr. Dirk Schubert (RadboudUMC, Nijmegen, The Netherlands) for supporting to run the experiments at their laboratory. Naoki Kogo was supported by a post-doctoral fellowship of Fonds voor Wetenschappelijk Onderzoek (FWO-Flanders post-doc grant 12L5115N, University of Leuven, 2014~2017) and a European Fellowship from Marie Curie Actions (794273, Radboud University, 2018~current). TN was supported by the EPSRC (EP/P006094/1) and European Union (785907). RvW was supported by grants by NWO-TTW (NESTOR and INTENSE).

## Supplemental Information

**Fig. S1.**
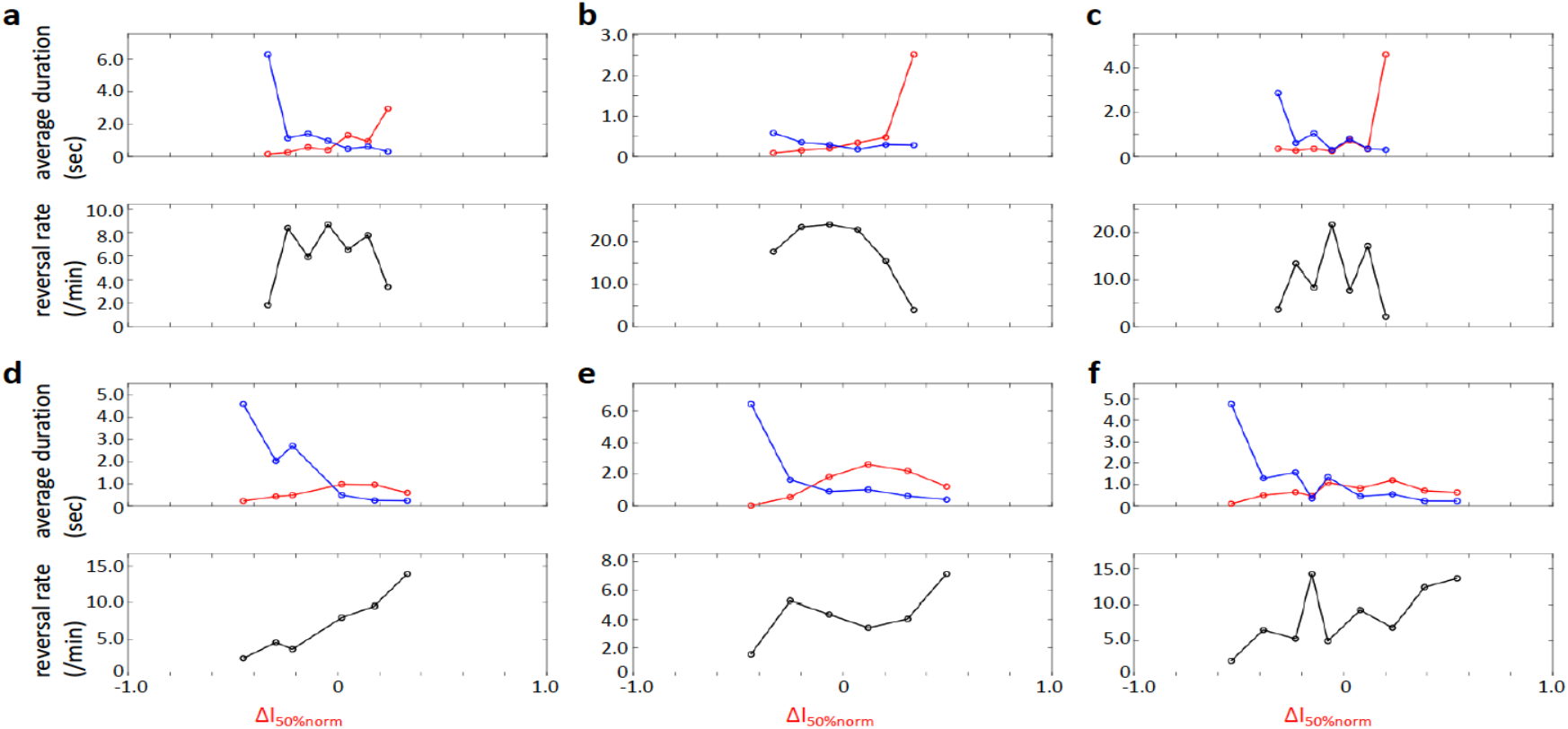
Examples of responses of average durations and reversal rate. They are shown (in non-normalized absolute values) as responses to the change of the depolarization current to one of the pair of pyramidal neurons (red) while the current to the other neurons was kept constant (blue). **a-c**: The example pairs where the reversal rates decreased when the current either increased or decreased from the control value (*I*_*50%*_). **d-f**: The examples where the reversal rates increased when the current increased from the control value (*I*_*50%*_). Note that, in the latter case, the increase of the average durations of the dominant neuron

